# Beyond Plug and Pray: Context Sensitivity and *in silico* Design of Artificial Neomycin Riboswitches

**DOI:** 10.1101/2020.06.18.159046

**Authors:** Christian Günzel, Felix Kühnl, Katharina Arnold, Sven Findeiß, Christina Weinberg, Peter F Stadler, Mario Mörl

**Affiliations:** Institute for Biochemistry, Leipzig University, Brüderstraße 34, D-04103 Leipzig, Germany; Bioinformatics Group, Institute of Computer Science, and Interdisciplinary Center for Bioinformatics, Leipzig University, Härtelstraße 16–18, D-04107 Leipzig, Germany; Max-Planck-Institute for Mathematics in the Sciences, Inselstraße Leipzig, D-04103 Leipzig, Germany; Institute for Theoretical Chemistry, University of Vienna, Währingerstraße 17, A-1090 Wien, Austria; Facultad de Ciencias, Universidad National de Colombia, Sede Bogotá, Colombia; Santa Fe Institute, 1399 Hyde Park Rd., Santa Fe, NM 87501, USA

**Keywords:** riboswitch design, synthetic riboswitch, regulatory RNA, neomycin aptamer, transcription regulation, leader hairpin, decoupling leader, context optimization, N1 aptamer

## Abstract

Gene regulation in prokaryotes often depends on RNA elements such as riboswitches or RNA thermometers located in the 5’ untranslated region of mRNA. Rearrangements of the RNA structure in response, e. g., to the binding of small molecules or ions control translational initiation or premature termination of transcription and thus mRNA expression. Such structural responses are amenable to computational modeling, making it possible to rationally design synthetic riboswitches for a given aptamer. Starting from an artificial aptamer, we construct the first synthetic transcriptional riboswitches that respond to the antibiotic neomycin. We show that the switching behavior *in vivo* critically depends not only on the sequence of the riboswitch itself, but also on its sequence context. We therefore developed *in silico* methods to predict the impact of the context, making it possible to adapt the design and to rescue non-functional riboswitches. We furthermore analyze the influence of 5’ hairpins with varying stability on neomycin riboswitch activity. Our data highlight the limitations of a simple plug-and-play approach in the design of complex genetic circuits and demonstrate that detailed computational models significantly simplify, improve, and automate the design of transcriptional circuits. Our design software is available under a free license on Github.^1^

## 1. Introduction

Riboswitches are *cis*-regulatory elements that modulate gene expression in response to small molecules or ions without direct involvement of proteins [1, 2]. They are typically embedded in the 5’ untranslated region (UTR) of messenger RNA (mRNA) and usually comprise two overlapping domains: a conserved *aptamer,* presenting the binding site for a specific ligand, and a genespecific *expression platform* that regulates the expression of the gene located downstream. Riboswitches can adopt mutually exclusive structures depending on the presence or absence of their ligand. The structural reorganization or stabilization of the aptamer upon ligand binding triggers a change of the structure of the expression platform [3]. The latter may affect either translation – e. g., by changing the accessibility of the ribosomal binding site – or transcription, e.g., by modulating an intrinsic terminator causing the premature dissociation of RNA polymerase (RNAP). The relatively small size of riboswitches facilitates their design and optimization [4], which makes them attractive for numerous applications in synthetic biology. The ability to construct orthogonal switches, for instance, makes it possible *in principle* to force a gene to respond to any cell-permeable and non-toxic ligand [5] and thus to create functional biosensors that turn on the expression of a certain reporter gene in response to both intracellular signals and environmental conditions. The practicality of such an approach, however, depends on how easily aptamers and expression platforms can be combined and embedded in an arbitrary sequence context without resorting to a labor-intensive trial-and-error procedure.

The antibiotic neomycin has a generic RNA binding affinity which allowed to engineer cognate aptamers by means of systematic evolution of ligands by exponential enrichment (SELEX) [6, 7]. The aptamer N1 was isolated by Weigand et al. [8] and shown to act as a translational roadblock in presence of neomycin in yeast. N1 binds to its ligand with an extraordinarily high affinity in the lower nanomolar range [9]. Its switching behavior in response to neomycin was analyzed *in vivo* in conjunction with various closing stems that modulate the expression of a green fluorescent protein (GFP) reporter located downstream. The shortest stem, designated M7, exhibited both the highest fluorescence intensity and the best ratio between its *on* and *off* state [9].

Based on the aptamer N1M7, we constructed a riboswitch that controls gene expression in *Escherichia coli* (E. coli). To this end, we focused on transcriptional regulation mediated by Rho-independent termination, a mechanism common in prokaryotes. It depends on a specific terminator structure, consisting of a GC-rich stem–loop followed by 7–9 uracil residues referred to as *poly-U stretch* [10, 11].

We used the *in silico* pipeline developed previously for designing theophylline and tetracycline riboswitches [12, 13]to integrate the N1M7 aptamer with artificial terminator hairpins. The designed neomycin riboswitches indeed increase the transcription level of the reporter mRNA in presence of neomycin *in vivo.* A detailed analysis of the constructs, however, shows that their function *in vivo* is context-dependent. In particular, we identify a 5’ leader hairpin as essential element for the function of these neomycin riboswitches. On the other hand, leader sequences may also abrogate the function by interfering with the riboswitch. We demonstrate here that folding simulations predict the interference and thus can be used to identify functional constructs *in silico.*

## 2. Materials and Methods

### 2.1. Cells

For cloning purposes, E. coli Top10 cells (Invitrogen) were used. Riboswitch-controlled expression of the enhanced green fluorescent protein (eGFP) was tested by *in vivo* fluorescence measurements using the neomycin-resistant E. coli SQ171, derived from E. coli MG1655 [14] (obtained from Kurt Fredrick, Columbus, OH).

### 2.2. Plasmid construction

As reporters, the enhanced green fluorescent protein gene *(egfp)* or the *β-*galactosidase gene (bgaB) were inserted into plasmid pRSF1030Tp [15] by restriction and ligation reactions (using the *Nco*I and *Xba*I site for egfp or *Age*I and *Kpn*I site for *bgaB*). Riboswitch constructs were synthesized and amplified by *de novo* PCRs. For *de novo* fragments longer than 90 bp, the construct was split in two or more overlapping fragments and extended by overlap extension PCR with flanking primers. The riboswitch sequences were inserted downstream of the *araBAD* promotor and upstream of egfp or bgaB through restriction and ligation reactions or mutagenesis PCR (for all LM constructs). All PCR reactions were conducted according to the protocol of HiFi DNA polymerase (PCR Biosystems). For overlap extension PCR, the flanking primers were added after 10 cycles, followed by amplification for additional 25 cycles. Plasmid isolations were performed according to the GeneJET Plasmid Miniprep Kit protocol (Thermo Scientific) for low-copy-number plasmids. All constructs were verified by sequencing.

### 2.3. *In vivo* measurements of fluorescence intensity

The *in vivo* eGFP fluorescence measurements were performed in E. coli SQ171. Precultures were incubated in 5 ml LB medium with 200 μg ml^−1^ trimethoprim over night at 37 °C and 200 rpm. 100 μl of the precultures were used to inoculate fresh cultures (5 ml), which were incubated at 37 °C and 200 rpm until an OD_600_ of 0.5. Expression of eGFP was induced by addition of L-(+)-arabinose (to 0.1% w/v) in the absence and presence of neomycin (250μg ml^−1^). After incubation for 3 h at 37 °C and 200 rpm, cultures were immediately stored on ice to measure the final OD_600_ and diluted with cold 1 × PBS to 1 ml (OD_600_ = 1.5). Samples were centrifuged at 8 000 rpm for 5 min, the pellet was washed twice with cold 1× PBS and resuspended in 500 μl 1× PBS (equaling OD_600_ = 3.0). eGFP fluorescence was measured from 503 nm to 600 nm with a fluorescence spectrometer (JASCO Deutschland GmbH, model: FP-8500, mode: emission, ex bandwidth: 2.5 nm, em bandwidth: 5nm, response: 1 s, sensitivity: low, ex wavelength: 488nm).

As a positive control, we combined a 5’ adenosine residue following the transcription start site (TSS) with the poly-U stretch of length eight and the terminator downstream region (TDR) of our riboswitch constructs. The poly-U stretch is known to promote translation [16] and, thus, makes the control comparable to the full riboswitch sequences.

### 2.4. RNA extraction and northern blots

Total RNA was isolated from E. coli SQ171 cells grown in the presence or absence of 275 μM neomycin until OD_600_ = 0.5 in 20 ml LB medium supplied with trimethoprim (200μg ml^−1^) and arabinose (0.1%). Total RNA was isolated using TRIzol™ (Invitrogen) according to the manufacturer’s protocol.

Northern blot analysis was performed as described by Köhrer and Raj-Bhandary [17] with modifications. 10μg of total RNA were mixed with 3× denaturing sample buffer (10mM Tris-HCl, pH 7.6, 80% formamide (v/v), 0.25% bromophenol blue and 0.25% xylene cyanol) and incubated for 1 min at 90 °C before electrophoretic separation on a 1% agarose gel containing 1× Tris-borate-EDTA (TBE) (89mM Tris, 89mM boric acid, 2mM EDTA, pH 8.0).

The gel (7 × 8 cm) was run at 70 V in 1× TBE for 1.5 h. RNA was transferred onto BrightStar™ Positively Charged Nylon Membranes (Ambion^®^) at 2 mA cm^−2^ in a semi-dry electroblotter (Thermo Fisher Scientific) for 45 min using 1 × TBE buffer. RNA fixation by cross-linking was performed by irradiation at 254 nm UV light for 60s (120 mJ cm^−2^ total) in a Stratalinker^®^ UV Crosslinker (Artisan Technology Group). The membrane was pre-hybridized for 14 h to 20 h at 37°C in 6× saline-saline sodium citrate buffer (SSC) containing 10× Denhardt’s solution and 0.5% SDS, followed by hybridization for 12h to 24h at 37°C (for bgaB) or 60°C (for *egfp*) in 6× SSC containing 0.1% SDS with 5’ radioactively labeled oligonucleotide probes for *bgaB* mRNA^2^, *egfp* mRNA^3^ or 5S ribosomal RNA (rRNA)^4^. Probes were radioactively labeled with *y*-^32^P-ATP (Hartmann analytic), unincorporated nucleotides were removed with NucAway™ Spin Columns (Invitrogen). The hybridized membrane was washed twice at room temperature with 6× SSC, 4× SSC and 2× SSC (each wash step 10 min). Signals were detected using a Typhoon phosphor imager (GE Healthcare Life Sciences, model: 9410). For quantitation of mRNA levels, northern blot signals of three independent colonies per construct (with and without neomycin) were analyzed using ImageQuant 8.1 (GE Healthcare Life Sciences). Membranes were stripped of radiolabelled probes by incubating in 0.5% SDS at room temperature for 1 h.

### 2.5. The N1M7 aptamer

The starting point for our *in silico* design is the sequence of the well-characterized neomycin aptamer N1M7, i.e., the aptamer N1 combined with the closing stem M7 consisting of five base pairs [8], cf. Fig. 1. The aptamer binds neomycin with a dissociation constant of *K_d_* = (9.2 ± 1.3) nM, corresponding to a stabilizing Gibbs free energy contribution of Δ*G* = (–11.4 ± 0.1) kcalmol^−1^ [9].

**Figure 1:**
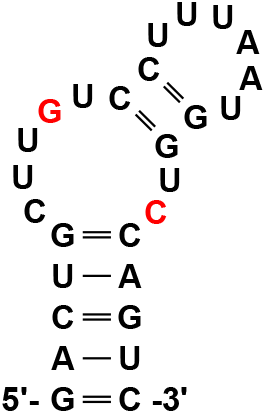
Sequence and secondary structure of the aptamer N1M7 [8]. Note that the minimum free energy structure predicted by the *ViennaRNA* package contains an additional GC pair at position 9 and 22 (colored red, energy difference ΔΔ*G* = – 0.7 kcal mol^−1^).

### 2.6. Probabilities of RNA conformations

RNA molecules can adopt a huge number of different secondary structures. In thermodynamic equilibrium, the probability to observe a particular structure *x* depends on its Gibbs free energy Δ*G*(*x*), which can be computed using the nearest-neighbor energy model [18], which specifies sequence-dependent stabilizing contributions for base pair stacking and destabilizing loop terms. The Boltzmann weight of *x* is defined as Z[*x*]:= exp(-Δ*G*(*x*)/(*RT*), where *R* is the universal gas constant and *T* is the absolute temperature. The *partition function Z*[*Y*] of a *set* of secondary structures *Y* is the sum of all the structures’ Boltzmann weights, i. e., Z[*Y*]:= ∑_*y∈Y*_ Z[*y*]. Writing Z:= Z[*X*] for the partition function of the full ensemble of the RNA, the equilibrium probability of a given structure *x* is Pr[*x*] = Z[*x*]/ Z, and, more generally, Pr[*Y*] = Z[*Y*]/ Z for a set of structures *Y*. Partition functions of the entire set of structures *X* as well as certain subsets *Y* (such as the set structures containing a specific hairpin loop, or the set of structures containing a particular base pair) can be computed by dynamic programming. We used the *ViennaRNA* package [19, 20]because it provides a versatile way to handle constraints [21] and thus to compute partition functions for a set *Y* of structures with desired structural features, e. g., a terminator hairpin or a binding pocket for a certain ligand.

### 2.7. Simulation of cotranscriptional folding

While initial folding intermediates of RNA form at time-scales of tens of microseconds, the formation of native hairpins appears at millisecond timescales [22, 23, 24, 25], and the refolding of secondary structure elements may take even longer. In comparison, RNA is transcribed by *E. coli* RNAP with a rate of 30nts^−1^ to 90nts^−1^ [26, 27]. Thus, RNA folding forms intermediate structures long before the entire molecule is transcribed, i. e., while only part of the transcript is available to form structures. The structures formed initially thus may refold as transcription proceeds. This process of cotranscriptional folding [28] plays an important role in particular for *transcriptional* riboswitches, since incomplete, metastable intermediate structures may be quite different from the thermodynamic ground state [29]. Cotranscriptional folding can be assessed either by stochastic sampling of folding trajectories [30, 31, 32] or by analyzing the energy landscapes for each elongation step [33]. We opted for the latter method.

For each length of the nascent transcript, we enumerated all secondary structures in an energy band above the ground state with *RNAsubopt* [34], also a component of the *ViennaRNA* package. Then we used *Barriers* [35] to produce a coarse-grained representation of the energy landscape comprising the low energy minima as well as the saddle points between them. *Barriers* assigns each structure to a basin of attraction. The individual landscapes were then integrated using *BarMap* [33]. In brief, *BarMap* determines the correspondence of energy basins in the landscapes of consecutive transcriptional elongation steps. This allows to efficiently simulate the folding dynamics. To determine the energy thresholds for the enumeration of structures with *RNAsubopt* in the individual landscapes, we used the quality scores for both the enumeration and the simulation provided by *BarMap-QA* [36]. We set a simulation stop time of 1 000 au (arbitrary time units) for *BarMap.* Simulation results were plotted with Grace [37]. Candidate riboswitches were screened to satisfy the following criteria: (i) the leader sequence does not interfere with the formation of the aptamer’s binding pocket, (ii) the binding pocket is significantly populated during the transcription of the spacer and the 5’ part of the terminator, and (iii) the terminator hairpin dominates the structure ensemble as soon as it is fully transcribed. For a precise description of the preparation steps, quality metrics, and the post-processing, we refer to [36]. To allow for easy reproduction, the entire simulation pipeline was packaged into a self-contained and publicly available Docker image.^5^

### 2.8. Design of transcriptional riboswitches

The design process consists of two separate phases. First, the riboswitch itself is designed from the given aptamer sequence. Second, for a set of designed riboswitch constructs, a decoupling leader (dL) is constructed, which isolates the riboswitches from the original leader (oL) sequence found upstream on the plasmid. This second step may become necessary if oL has the potential to interfere with the structure formation of the designed riboswitches (cf. Section 3.1). The full process is depicted as a flowchart in Fig. 2.

**Figure 2:**
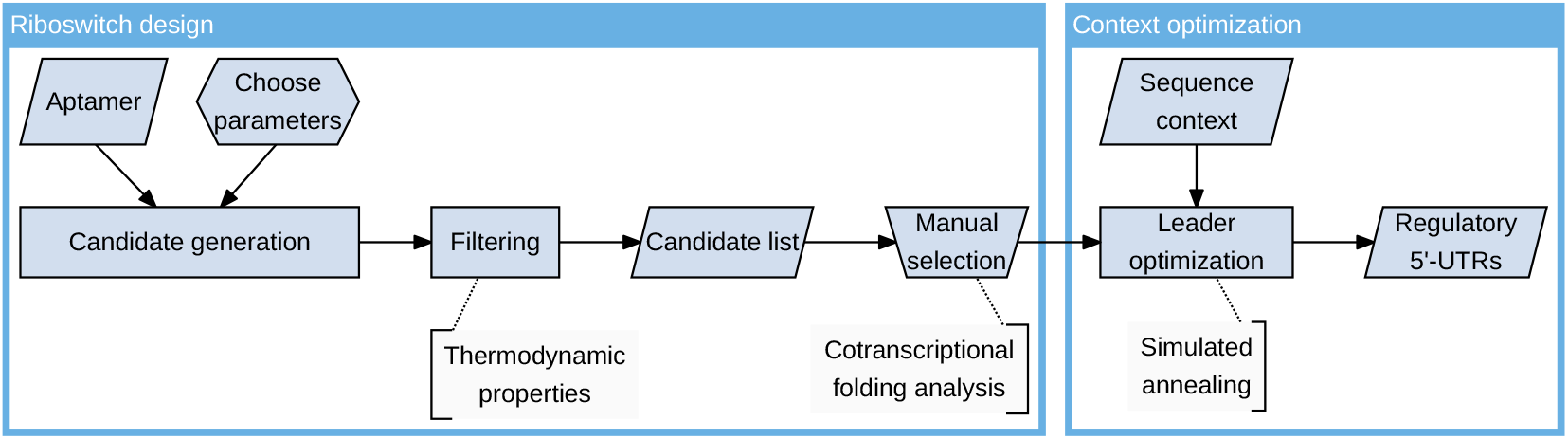
Flowchart of the design process used to create neomycindependent riboswitches and to include them into regulatory 5’ UTRs. Starting from a given aptamer sequence and structure, riboswitch candidates are generated and selected as described in the text.

To engineer the switch, a software pipeline similar to the one described by Wachsmuth et al. [12] including updated filter rules has been used. Briefly, this software is given the aptamer sequence and its ligand-binding structure as input, generates riboswitch candidates according to a specified pattern *(generation step),* and then applies a set of filters to remove unsuitable candidates *(filtering step*).

The generation step appends a random spacer sequence of varying length to the aptamer sequence. Then, the reverse complement of the last *k* nucleotides of the aptamer sequence is appended, forming a stable hairpin loop with the aptamer and disrupting the formation of the ligand-binding structure. The value of *k* varies from two to half of the aptamer length among the candidates. In addition, a U-stretch consisting of eight uracil residues is added, completing the structure of an intrinsic terminator [38].

To remove potentially faulty constructs, the filtering step applies a set of filters as described in Wachsmuth et al. [12], removing each sequence that violates any of the filter rules. The rules ensure correct terminator formation in the minimum free energy (MFE) structure and scan for possibly interfering intermediate structures by the means of thermodynamic folding simulations of a set of subsequences of the full switch. Additionally, we added a probability-based filter ensuring that, for the full sequence, the terminator structure forms with a probability of at least 95%. Also, it was ensured that at least two base pairs of the terminator hairpin can form despite the presence of the binding-competent aptamer structure. These *seed base pairs* facilitate the rapid formation of the terminator hairpin when the ligand is not present since the zippering of a helix happens at a higher rate than the nucleation of the first base pair [22, 23]. Finally, promising candidates resulting from this selection process where analyzed using cotranscriptional folding simulations as described in the previous section.

### 2.9. Designing a decoupling leader sequence

To prevent the predicted interference of the original leader located upstream in the 5’ UTR with the designed riboswitches, a sequence insert that effectively decouples leader and riboswitch has been designed. To this end, we defined an objective function *F* as described below and optimized a sequence with respect to *F*. The sequence with the best score was then cloned into the vector immediately upstream of the riboswitch, cf. Fig. A.1 in Appendix A. To keep the required experimental effort as low as possible, only one optimized leader suitable for all riboswitch constructs was designed.

Let *ℓ* denote the original leader, and *R* = {*r*_1_,…, *r_n_*} be a set of *n* riboswitches. The goal is to construct an insert *x_m_* ∈ {*A, U, G, C*}^*m*^ of length *m* that minimizes the objective function *F*(*x_m_* | *ℓ, R*) measuring the interference of the upstream sequence *ℓx_m_* with each of the riboswitches, subject to a constant length *m*. A natural measure for the isolation of *ℓx_m_* and the riboswitches is the probability

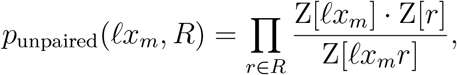

that all base pairs occur either in the upstream part *ℓx_m_* of the sequence (structures in Z[*£x_m_*]) or within the riboswitches (structures in Z[*r*]), but not between the two substructures. Using Δ*G*(*x*) = –*RT*lnZ[*x*], it can be expressed equivalently in terms of Gibbs free energies, with the added benefit of numerical stability. We therefore use the following objective function:

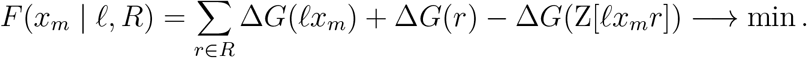

The optimization of *x_m_* was carried out using a standard simulated annealing procedure [39], starting at a random sequence and applying single nucleotide mutations to the insert to generate new candidates. A proposal sequence 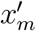 was *always* accepted if it performed better than the current state *x_m_* (i. e., if 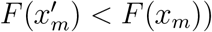, and otherwise with a probability of 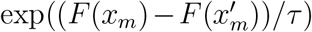, where the “annealing temperature” *τ* was slowly decreased with time. More specifically, we set the initial temperature to *τ*_0_ = 1000 and, after each mutation, cooled it down by letting *τ*_*n*+1_ = 0.97 · *τ_n_*. We used the rejection of 3*m* consecutive proposals as stopping criterion, where 3*m* is the number of neighbor sequences which can be obtained by applying a single point mutation to the candidate insert of length *m*. This ensures that, on average, about two thirds of its neighbors are sampled before terminating the optimization run. During the optimization, *m* was fixed because longer sequences generally have a higher potential to form stable structures fulfilling the objective, which could cause degenerate optimization runs with candidates of ever-growing sequence length.

## 3. Results

Many of our results are presented as bar plots of fluorescence intensity. The numerical values and raw data used to generate these can be found in Appendix B, Table B.1. Sequence data is available as a supplementary spreadsheet file.

### 3.1. Design of artificial neomycin riboswitches

Using computational predictions of ligand-induced differences in RNA secondary structure formation, we designed a set of transcription-regulating riboswitches based on the well-characterized neomycin aptamer N1M7, Fig. 1. These were evaluated *in vivo* in E. coli strain SQ171 using eGFP as reporter. This strain is neomycin-resistant, as all endogenous rrn operons were deleted and replaced by a plasmid-borne version carrying the mutated neomycin target site A1408G [14].

The plasmid used for assaying our riboswitch constructs contained a leader sequence we termed *original leader (oL)* immediately downstream of the promoter. Since its effect on transcription was unknown, we did not attempt to delete it. *In silico* analysis of the constructs suggested, however, that the oL sequence interferes with the structure formation of the N1M7 aptamer and thereby prevents the correct folding of its ligand binding pocket, Fig. 3. To remedy this issue, we designed a decoupling leader (dL) with a length of 15 nt that reduces the probability that base pairs form between oL and the riboswitch domain to less than 1%. It does so by forming the stable hairpin LH1, i.e., LH1 = oL + dL (cf. Fig. A.1).

**Figure 3:**
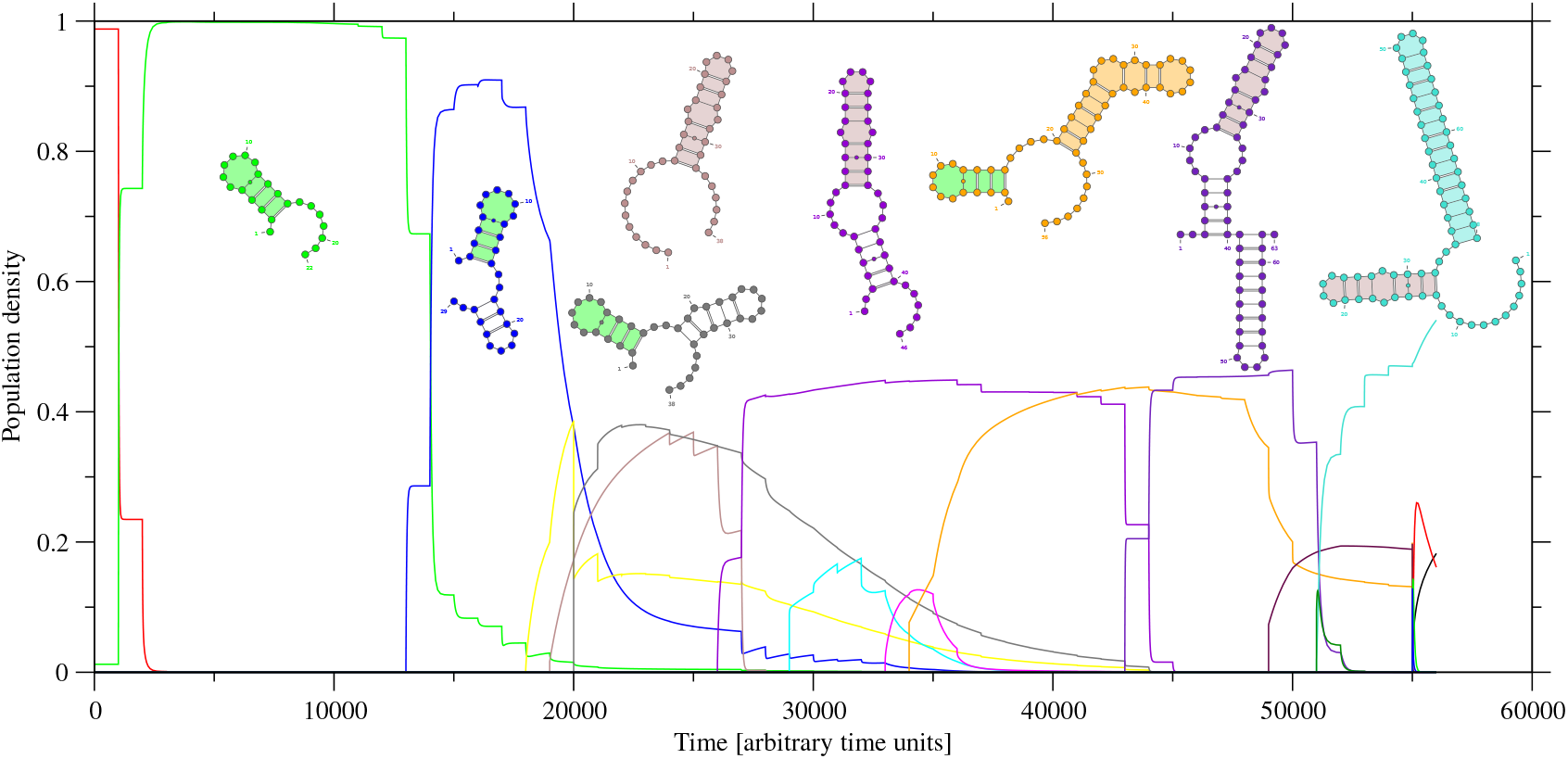
Influence of 5’ the original leader (oL), which is predicted to interfere with the used aptamer *in silico*, on the kinetics of riboswitch N1M7-D. The individual curves show the population of different macrostates during the elongation of the transcript. The most stable structure of highly populated states is shown in the respective color. Important sub-structures are shaded.

Four candidates prefixed with LH1 were selected for in-depth analysis, Fig. A.2. Fluorescence measurements of reporter gene expression (cf. Fig. 4A-B) showed that LH1-N1M7-C and LH1-N1M7-D function as *on*-switches, i.e., the ligand causes up-regulation of the reporter. In contrast, LH1-N1M7-A remains in a permanent *off* state while LH1-N1M7-B exhibits a permanent *on* state. As a representative example, northern blots in the presence and absence of neomycin were used to validate that LH1-N1M7-D regulates the amount of transcripts in the cell, Fig. 4C-E. Blotting experiments targeting egfp mRNA resulted in a low signal-to-noise ratio. Hence, we replaced the reporter gene by the well-established *β*-galactosidase (bgaB) reporter that we have used successfully for detection in northern blot analyses of theophylline-dependent riboswitches [12].

**Figure 4:**
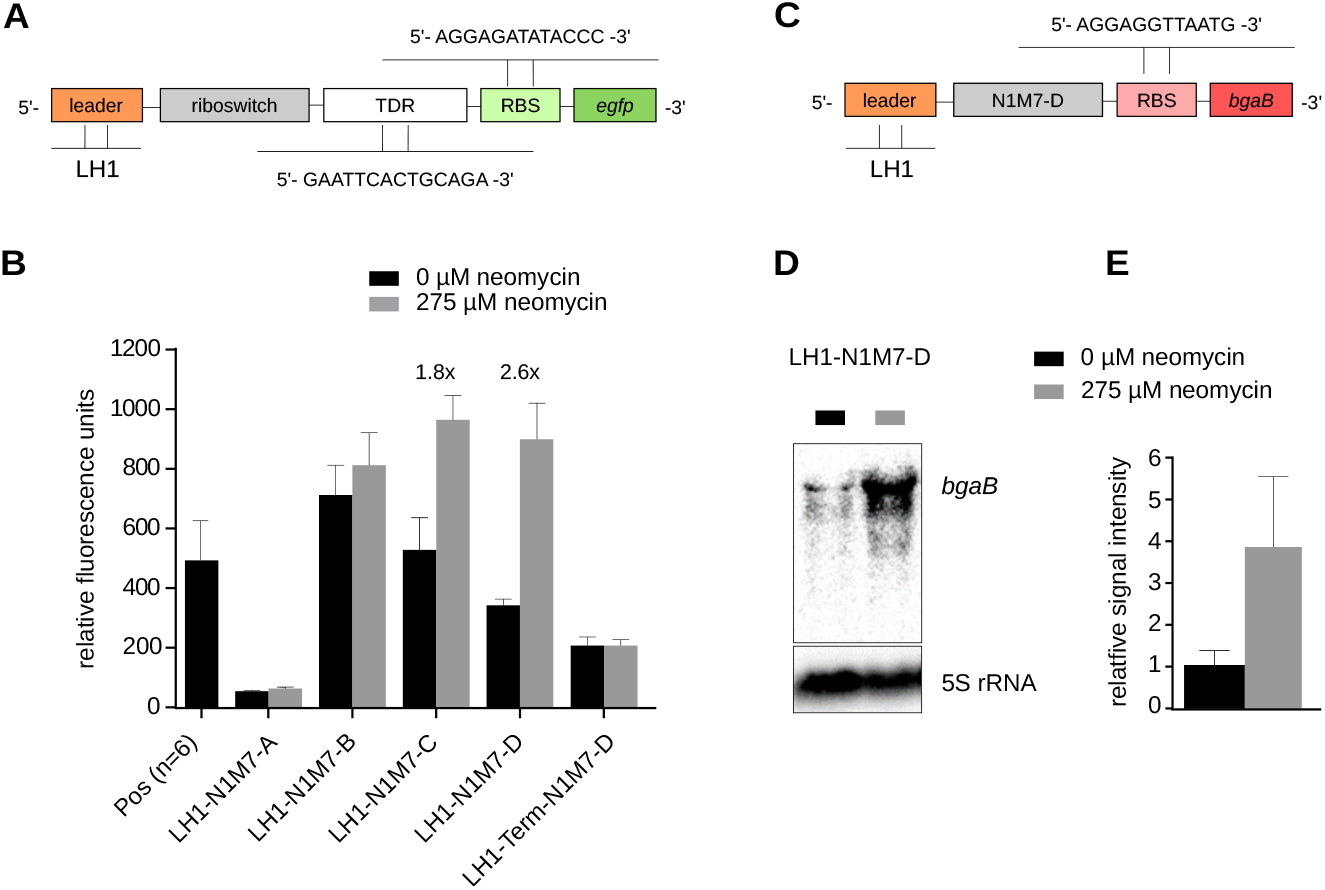
Design and analysis of N1M7 riboswitch constructs. (*A*) Schematic overview of the riboswitch constructs for the eGFP assay. The riboswitches are flanked upstream by LH1 and downstream by the terminator downstream region (TDR), ribosomal binding site (RBS) and the enhanced green fluorescent protein gene (*egfp*). *(B*) Fluorescence intensity of the measured N1M7 constructs in the absence (black) and presence (grey) of neomycin. The positive control, a transcript consisting of a single adenosine residue followed by a poly-U stretch and the TDR, shows the fluorescence intensity without leader and riboswitch sequence. The terminator efficiency was verified with construct LH1-Term-N1M7-D. Here, the 5’ part of the aptamer sequence that is not overlapping with the intrinsic terminator was deleted, such that the terminator forms irrespective of the presence of neomycin. If not indicated otherwise, measurements were performed using three independent replicates. *(C*) Schematic presentation of a neomycin riboswitch construct for northern blot analysis. Downstream of the riboswitch, the construct carries the *β*-galactosidase reporter gene (*bgaB*) including the RBS published in Wachsmuth et al. [12]. Due to the design strategy, this construct does not carry a TDR. (*D*) Northern blot with 10 μg of total RNA of E. coli strain SQ171 per lane regulated by LH1-N1M7-D, in the absence and presence of neomycin. 5S rRNA was used as internal standard. (*E*) Northern blot quantification of *bgaB* expression of three independent samples.

In essence, oL forms two distinct, stable structures: a small hairpin of four base pairs (green) that is compatible with the aptamer’s binding pocket (orange), and a larger hairpin (brown) incorporating an interior loop that is extended later (light and dark purple) and interferes with the aptamer region. Both structural shapes are equally populated with about 40%, which means that the aptamer is not available for binding the ligand in about every other transcribed molecule. The terminator (cyan) forms immediately after it has been transcribed. This explains the permanent *off* state that has been observed for Lpl-N1M7-D experimentally, cf. Fig. A.4.

### 3.2. Riboswitch N1M7-D requires a 5’ hairpin structure

Instead of designing a decoupling sequence for oL with a complicated optimization method as described in the previous section, the more obvious approach seems to simply remove oL from the plasmid and place the riboswitch candidates directly downstream of the TSS represented by an adenosine residue. It turns out, however, that the removal of all leader sequences from the functional riboswitch LH1-N1M7-D yields a *non-functional* construct termed N1M7-D, which is no longer sensitive to neomycin (Fig. 5C). The riboswitch function was rescued by inserting a second, computationally designed leader hairpin LH2 (Fig. 5A) resulting in the functional riboswitch LH2-N1M7-D, Fig. 5C.

**Figure 5:**
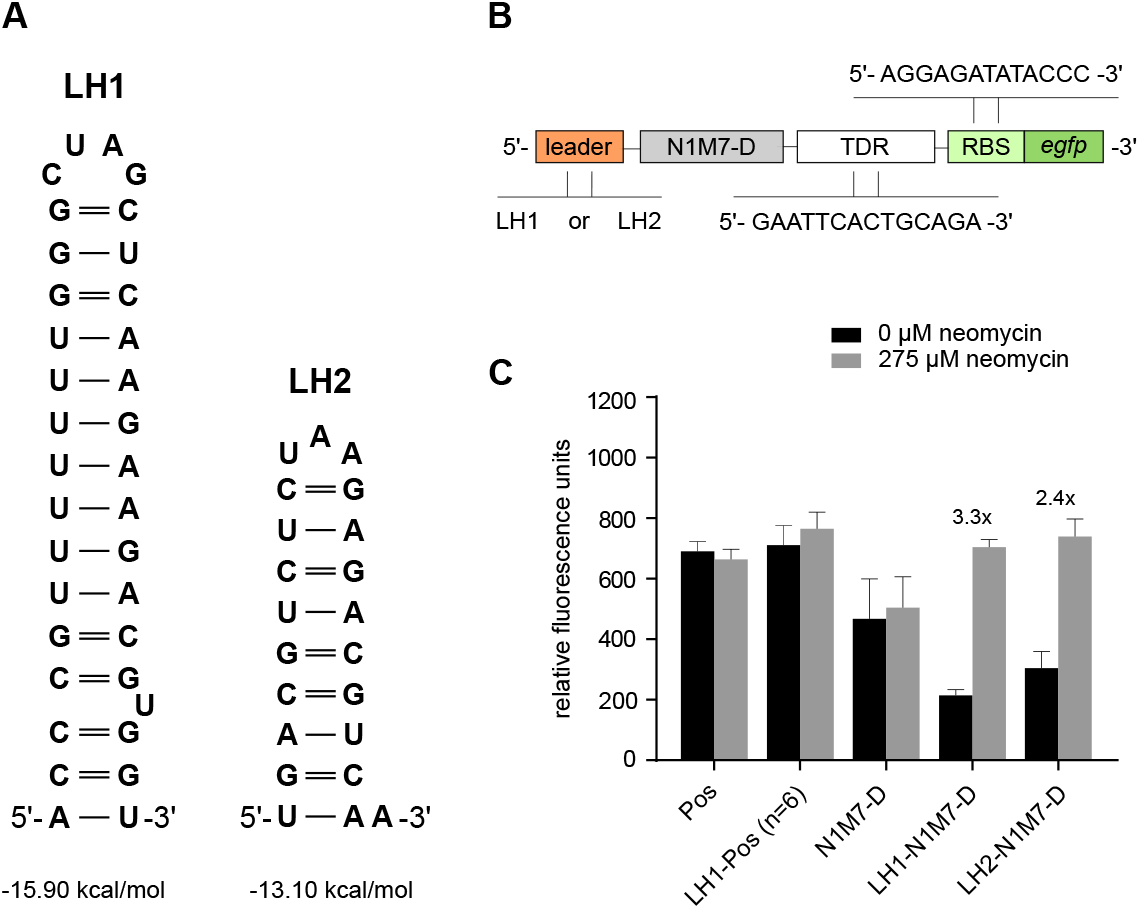
Fluorescence intensity of the riboswitch N1M7-D in conjunction with two different leader hairpins LH1 and LH2. (*A*) Sequence and secondary structure of LH1 and LH2. (*B*) Schematic overview of the constructs used in panel C. (*C*) Fluorescence intensity of the two leaders LH1 and LH2 placed upstream of N1M7-D. Legend as described in Fig. 4.

To rule out that the rescue is the consequence of a changed TSS, or a direct interaction of the leader hairpin with the riboswitch domain, we tested two short unstructured leader sequences U1 and U2, 12 nt and 14 nt in length, Fig. 6A. In the computational design of U1 and U2, base pairing interactions with the riboswitch domain were avoided. Both constructs, U1-N1M7-D and U2-N1M7-D, were functional with a fold change of 1.6 and 1.4 (respectively), however, their fluorescence intensity was significantly reduced, Fig. 6C. As with N1M7-D, it resembled the *off* state of the functional switch. Next, we re-added the leader hairpin LH1 to the 5’ ends of these impaired riboswitches. As a result, the original fluorescence activity was restored for both LH1-U1-N1M7-D and LH1-U2-N1M7-D, Fig. 6B.

**Figure 6:**
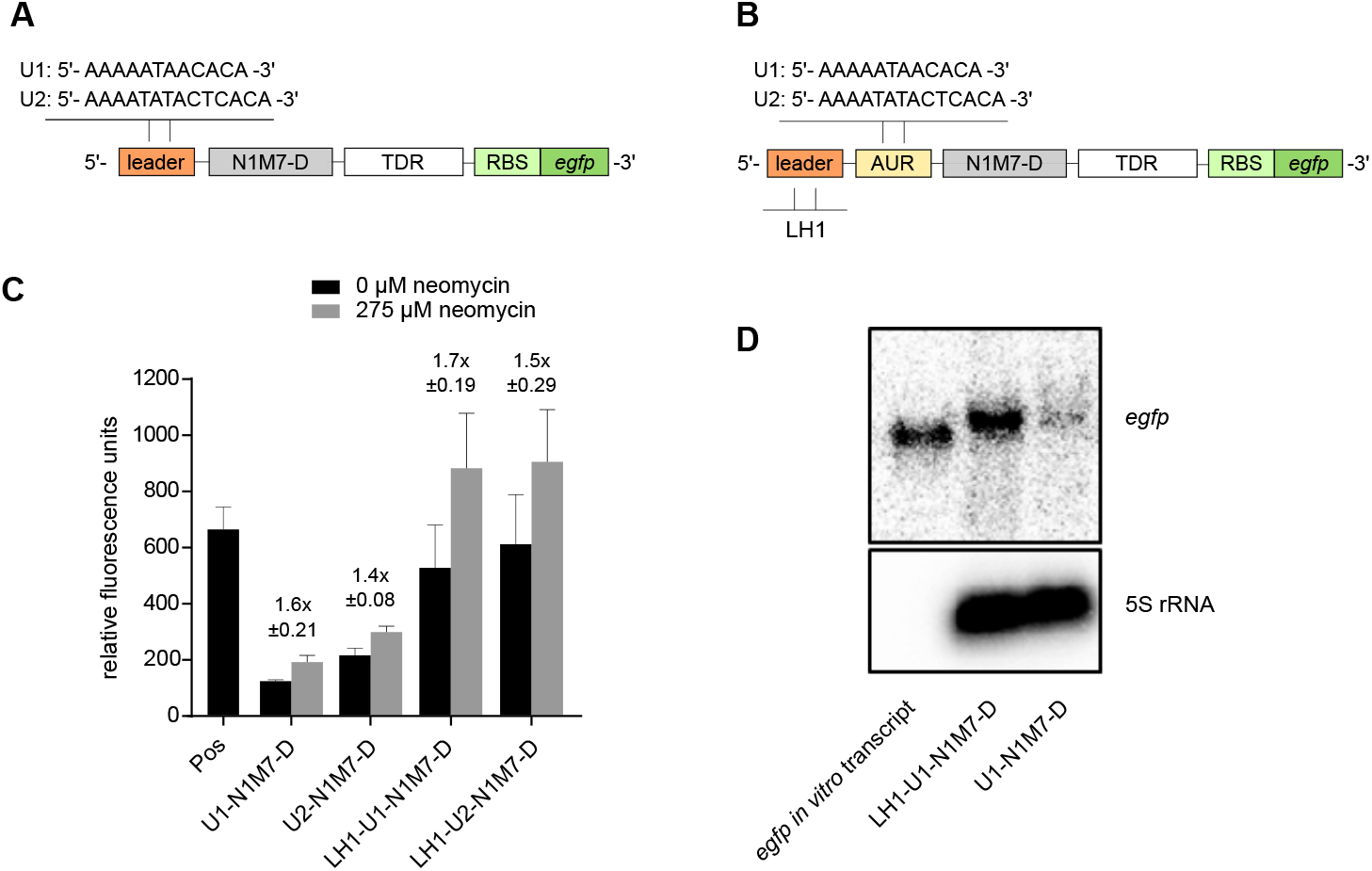
Fluorescence intensity of the N1M7-D riboswitch with short unstructured leaders. (*A*) Schematics showing the position of the unstructured sequences U1 and U2 as leaders upstream of N1M7-D and *(B*) as aptamer upstream region (AUR) between LH1 and N1M7-D. *(C*) Fluorescence intensity of the constructs shown in (*A*) and (*B*). Legend as in Fig. 4. *(D*) Northern blot of 10 μg of total RNA per lane from E. coli strain SQ171 regulated by U1-N1M7-D or LH1-U1-N1M7-D, in the presence of 275 μM neomycin. Herein, *egfp* was used as reporter gene and 5S rRNA as internal standard. A 1100 nucleotide (nt) *egfp in vitro* transcript was used as positive control. Note that the band of *in vivo egfp* is located about 100-200 nt higher due to the additional rrnB T1 and T2 terminator length, which was not a factor in size calculation for the *in vitro* transcribed control.

### 3.3. Destabilizing the leader hairpin decreases reporter gene expression

Since unstructured leader regions abrogated the switching behavior, we investigated whether the stability of the leader hairpin has a systematic effect on the riboswitch. To this end, we first introduced destabilizing point mutations into the leader hairpin LH1. We obtained two new leaders LM1 and LM2 (Fig. 7A), each lacking two base pairs. The destabilization was reverted by compensatory mutations in LM1C and LM2C, restoring the stem of LH1. We found that none of these mutations significantly affected the functionality of the riboswitch N1M7-D. The activation ratios of these constructs range between 2.8-fold to 3.4-fold, comparable to LH1-N1M7-D, cf. Table B.2.

**Figure 7:**
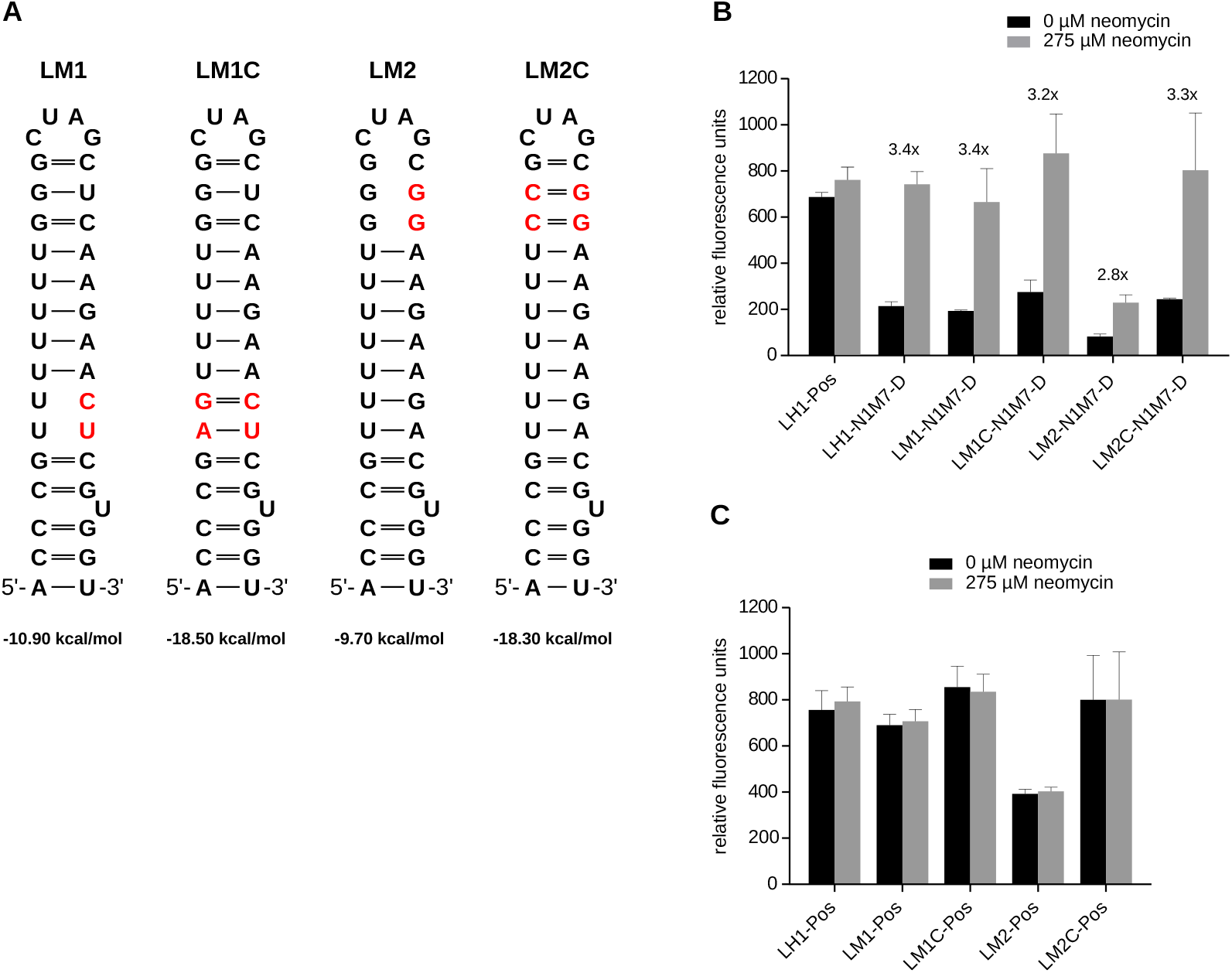
Impact of point mutations in the leader hairpin LH1. Fluorescence intensity of the modified leader LM1, LM1C, LM2 and LM2C (*A*) in conjunction with N1M7-D (*B*) or Pos (*C*), along with the controls LH1-N1M7-D or LH1-Pos. Legend as described in Fig. 4. Mutation details and minimum free energies are given in Table B.2.

There is, however, a trend in the overall fluorescence intensity: most stable hairpins lead to a higher eGFP expression, both in presence and absence of neomycin (Fig. 7B). Constructs lacking the riboswitch domain, i.e., those consisting of a leader hairpin and the positive control (Fig. 7C), show the same trend in the overall fluorescence intensity.

Similarly, we destabilised the alternative leader hairpin LH2, obtaining hairpins LM3, LM4 and LM5, Fig. A.3A. The construct LM3-N1M7-D, containing the least stable leader hairpin with an MFE of Δ*G* = −5.5 kcal mol^−1^, exhibits a constitutive *on* state (Fig. A.3B) but still shows a moderate acti-vation rate of about 1.5. Two compensatory mutations restore the hairpin (LM3C, Δ*G* = –13.9 kcal mol^−1^) to almost the same folding energy as LH2 (Δ*G* = –13.1 kcalmol^−1^) and rescue the function of the riboswitch. LM4-N1M7-D, with Δ*G* = –10.1 kcalmol^−1^, shows almost no difference in the activation rate or fluorescence intensity compared to LH2-N1M7-D or LM3C-N1M7-D. The fluorescence intensity of LM5-N1M7-D was decreased compared to LM4-N1M7-D, while the activation ratio virtually remained unchanged.

In summary, we observe that (i) a sufficiently stable leader hairpin appears to be required for functional constructs and (ii) the stability of the leader hairpin correlates with the constructs’ fluorescence intensity.

### 3.4. Upstream sequence context may impair riboswitch function

As mentioned before, we predicted that oL interferes with the aptamer N1M7 using *in silico* simulations. To verify this finding experimentally, we inserted oL between the leader hairpins and the riboswitch domains, resulting in the constructs LH1-oL-N1M7-D and LH2-oL-N1M7-D. In addition, we analysed oL-N1M7-D without any leader hairpin. All three constructs showed no neomycin-dependent regulation of eGFP expression *in vivo,* cf. Fig. A.4, even though the corresponding constructs without oL between LH1 and N1M7-D (namely LH1-N1M7-D, LH1-U1-N1M7-D, and LH1-U2-N1M7-D) were functional, cf. Section 3.2.

The expression level of LH1-oL-N1M7-D or LH2-oL-N1M7-D resembles the *off* state of the functional switch LH1-N1M7-D with a fluorescence intensity of 200–400 RFU (relative fluorescence units). Without a leader hairpin upstream of oL, the fluorescence reached an intensity of about 150 RFU. These low fluorescence intensities combined with the disturbed switching behaviour is in accordance with the predicted interaction of the oL with the neomycin binding pocket of the aptamer sequence N1M7. According to the simulations, this interaction not only prevents the formation of the neomycin binding pocket, but also facilitates the emergence of the terminator hairpin.

## 4. Discussion

In this work, we combined a synthetic neomycin aptamer with computationally designed terminator hairpins that abrogate transcription in the absence of the aptamer-specific ligand. The design process followed our earlier successful construction of theophylline- and tetracycline-dependent, transcription-regulating riboswitches [12, 13]. Here, we employed the neomycin aptamer N1M7, exhibiting an extraordinarily high binding affinity of *K_d_* = (9.2 ± 1.3) nM [9], as ligand sensor. Of the four computationally designed constructs that were tested *in vivo*, LH1-N1M7-C and LH1-N1M7-D were functional neomycin riboswitches. Northern blot analysis proved that LH1-N1M7-D acts as a transcriptional regulator, as intended. This demonstrates that our *in silico* approach can be successfully applied to novel aptamers.

The function of our artificial neomycin riboswitches depends on a stable 5’ hairpin structure. A plausible explanation for this requirement is an increased resistance to mRNA degradation: the half-life of a mutant *ompA* transcript increases 3-4-fold when such a hairpin is added upstream of its single-stranded 5’ UTR sequence [40]. ROSE elements – RNA thermometers in the 5’ UTR controlling many small heat shock genes in E. coli – consist of at least two consecutive hairpins, and only the last one is thermosensitive and regulating the mRNA‘s translation rate [41]. In gram-negative bacteria including *E. coli*, mRNA degradation is carried out by the degradosome, a complex of several proteins including Ribonuclease E (RNase E), supported by RNA 5’ pyrophosphohydrolase (RppH) [42, 43]. Stable 5’ structures are known to effectively obstruct the enzymatic activity of RppH and, thus, of the degradosome, explaining mRNA stabilization [44]. Our data show that the effect of the 5’ hairpin depends on its thermodynamic stability rather than its sequence, lending support to the hypothesis that stable 5’ hairpins play a general protective role in the process of mRNA degradation. An increased life-time of the mRNA would also explains the observed correlation between thermodynamic stability and fluorescence intensity. In the positive control, we observed no dependence of fluorescence intensity on the leader hairpin. However, this construct forms a very stable hairpin (instead of the flexible aptamer structure), which likely explains its stability.

While the leader hairpin is crucial for our neomycin riboswitch, no corresponding structure was required for similar theophylline or tetracycline riboswitches [12, 13], cf. Fig. A.5 for an overview of their thermodynamic ground states. We suspect that this is the consequence of differences in aptamer stability. While the neomycin aptamer has an MFE of Δ*G* = –6.2 kcal mol^−1^ [9], both the theophylline aptamer used by Wachsmuth et al. [12] (Δ*G* = –12.1 kcalmol^−1^) and the tetracycline aptamer used by Domin et al. [13] (Δ*G* = –19.8 kcal mol^−1^) are considerably more stable. We therefore hypothesize that, in the presence of the ligand, a highly stable aptamer-ligand complex sufficiently protects the transcript from degradation. In absence of the ligand, the terminator hairpin forms rapidly and leaves the transcript with an unstructured 5’ end, facilitating its quick degradation and thus contributing to a high activation ratio. The observed change in fluorescence activity upon ligand binding is therefore a result of two independent effects: an increased transcription rate due to the suppression of the terminator hairpin formation, and a reduced transcript degradation due to the stabilization of the aptamer structure located at its 5’ end.

Synthetic biology is concerned with solving sophisticated design problems with minimal expenditure of time, effort and money and thus strives to construct complex genetic circuits as a combination of fully modular, independently optimized components. Natural biological systems, however, have not evolved to adhere to the paradigm of engineering. It is well-documented, therefore, that the embedding of engineered circuits into living organisms faces a multitude of problems, mostly from unintended and often unexpected interaction with the natural context. Orthogonal systems attempt to avoid or at least minimize this problem [45]. Some quite spectacular success stories, e.g., the construction of re-engineered versions of secondary metabolite biosynthetic pathways [46] or the assembly of artificial operons [47] have raised the hope for a “plug-and-play” synthetic biology that allows the usage of pre-fabricated components in a combinatorial fashion. Even within an orthogonal system, however, it appears that this goal will remain elusive in many cases, at least in the strict sense of devising context-independent components [48]

Our data show that already conceptually very simple devices such as transcriptional riboswitches are more than a simple concatenation of building blocks. This is not surprising, given that the function of a transcriptional riboswitch is not just determined by the properties of aptamer and terminator, but depends on a delicate balance of mutually exclusive structural features and a carefully orchestrated time course of folding and refolding during its transcription [49]. As a consequence, prefabricated modules require adaptation and partial redesign to properly function in combination. In the case of RNA devices, the intrinsically global nature of RNA structure formation, in which base pairs are not restricted to local interaction, explains the imperfect modularity. At the same time, computational models of RNA folding are capable of capturing the undesired side effects and make it possible to algorithmically optimize novel designs based on existing components.

In previous work, for example, we observed that only one of two terminator hairpins functioned properly in the synthetic theophylline riboswitch RS10 [50]. Using a cotranscriptional folding algorithm, we found that the inactive construct harbored an intermediate structure that likely acted as a kinetic trap, delaying the formation of the terminator hairpin sufficiently to render it inactive.The adaptation of the interaction between the modular components is, by itself, not necessarily sufficient, as the presented work shows. The neomycin switches depend on an obligatory 5’ leader hairpin, presumably because the riboswitch itself is not sufficiently stable in itself to protect the mRNA from degradation. In addition, it needs to be adapted to the sequence context of the transcript to avoid interference of the leader sequence with the riboswitch domain – again an effect that can be captured by the RNA secondary structure model and remedied by adapting the constructs *in silico*.

Taken together, we conclude that complex RNA devices cannot be engineered by simply combining pre-optimized components in a plug-and-play fashion. Modular components can be combined and put into a functional context, however, with the help of *in silico* simulation techniques. We can therefore do better than “plug and pray”: computationally, we can adapt the modules to function in context and optimize the constructs as a whole. We are confident that the reliability of the computational predictions will continue to improve as our understanding of the individual components and their underlying mechanisms of action evolves.

## Supporting information

Sequence data for all constructs

Raw eGFP assay data

## Acknowledgements

We thank Julia E. Weigand and Beatrix Süß for their support and helpful discussions concerning the neomycin aptamer N1M7. Furthermore, we thank Kurt Fredrick for the neomycin-resistant *E. coli* strain SQ171 and John E. Cronan for the Plasmid pRSF1030Tp.

This work was supported in part by the German Research Foundation (DFG; grants STA 850/15-2 and MO 634/9-2).

## List of Abbreviations

This work makes use of the following abbreviations:

au: arbitrary time units
AUR: aptamer upstream region
bgaB: *β*-galactosidase
*bgaB*: the *β*-galactosidase gene
dL: decoupling leader
*E. coli*: *Escherichia coli*
eGFP: the enhanced green fluorescent protein
*egfp*: the enhanced green fluorescent protein gene
GFP: green fluorescent protein
LH: leader hairpin
LM: mutated leader
MFE: minimum free energy
mRNA: messenger RNA
oL: original leader
Pos: positive control
RBS: ribosomal binding site
RFU: relative fluorescence unit
RNA: ribonucleic acid
RNAP: RNA polymerase
RNase E: Ribonuclease E
RppH: RNA 5’ pyrophosphohydrolase
rRNA: ribosomal RNA
SELEX: systematic evolution of ligands by exponential enrichment
SSC: saline sodium citrate buffer
TBE: Tris-borate-EDTA
TDR: terminator downstream region
TSS: transcription start site
U: unstructured region
UTR: untranslated region

## A. Supplementary Figures

**Figure A.1:**
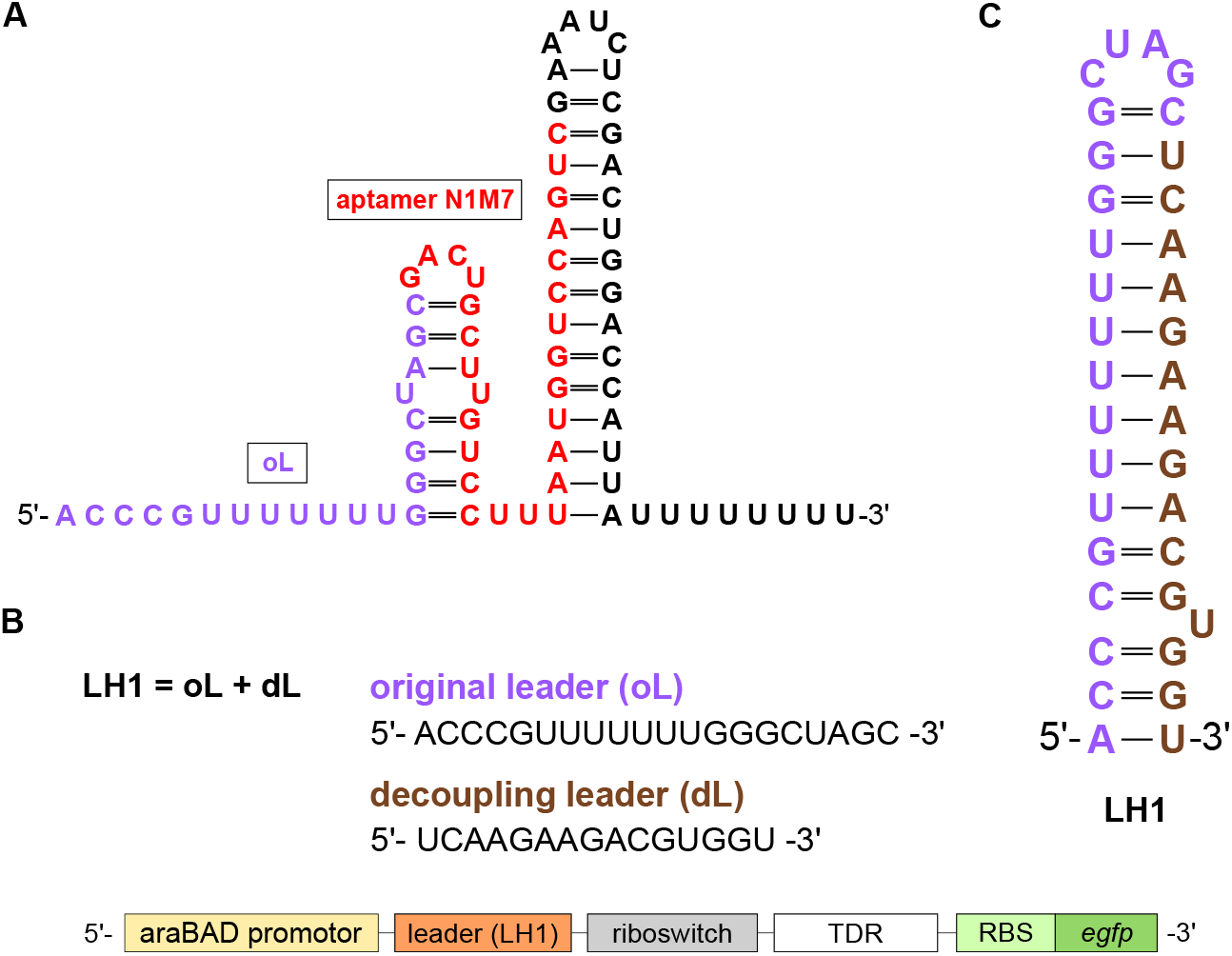
A decoupling leader for riboswitches. (*A*) Secondary structure of riboswitch N1M7-D with the original leader (oL) as computed by free energy minimization [20]. (*B*) Schematic of riboswitch-surrounding elements in the plasmid pRSF1030Tp-SD-eGFP. A decoupling leader was inserted downstream of oL to protect the riboswitch from leader-dependent misfolding by forming the stable hairpin LH1 depicted in (*C*).

**Figure A.2:**
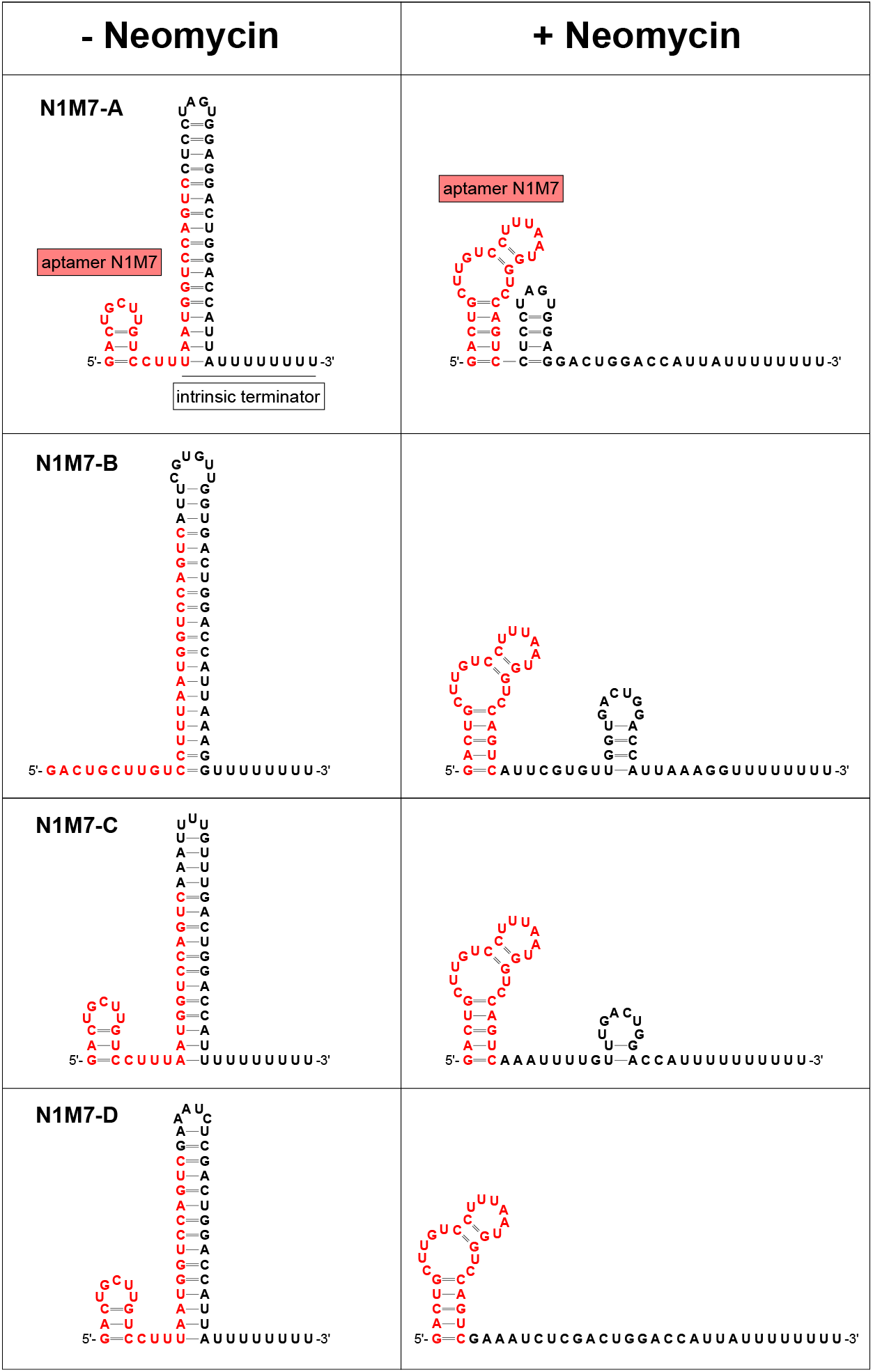
Secondary structure of the riboswitch constructs with the aptamer N1M7 used for the eGFP assay in Fig. 4.

**Figure A.3:**
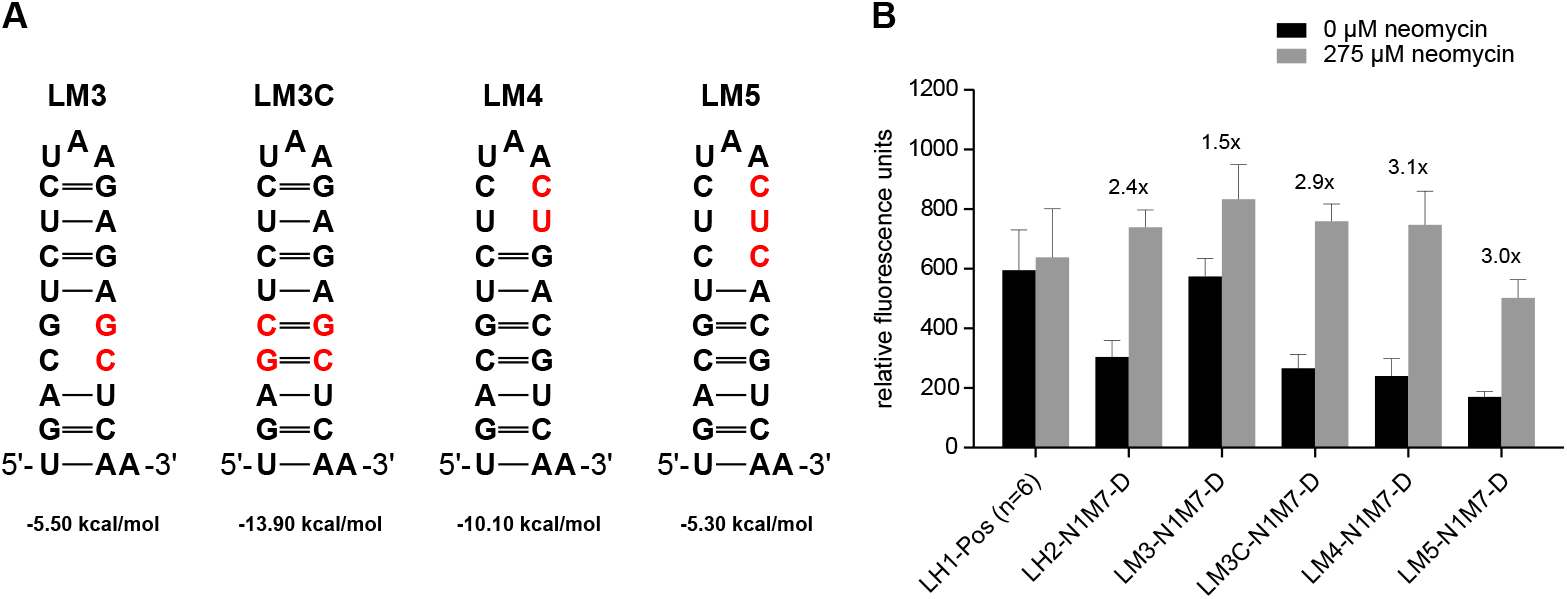
Impact of point mutations in the leader hairpin LH2. Fluorescence intensity of the modified leaders LM3, LM3C, LM4 and LM5, folded into their predicted secondary structure (*A*), in conjunction with N1M7-D (*B*). LH2-N1M7-D and LH1-Pos were used as control. Legend as described in Fig. 4.

**Figure A.4:**
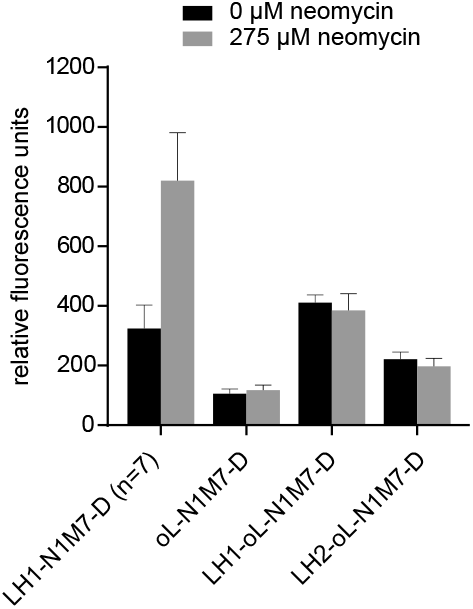
Influence of a sequence predicted to interfere with N1M7-D by *in silico* methods. The fluorescence intensity of N1M7-D with original leader (oL) in conjunction with LH1 or LH2 is shown. Legend as described in Fig. 4.

**Figure A.5:**
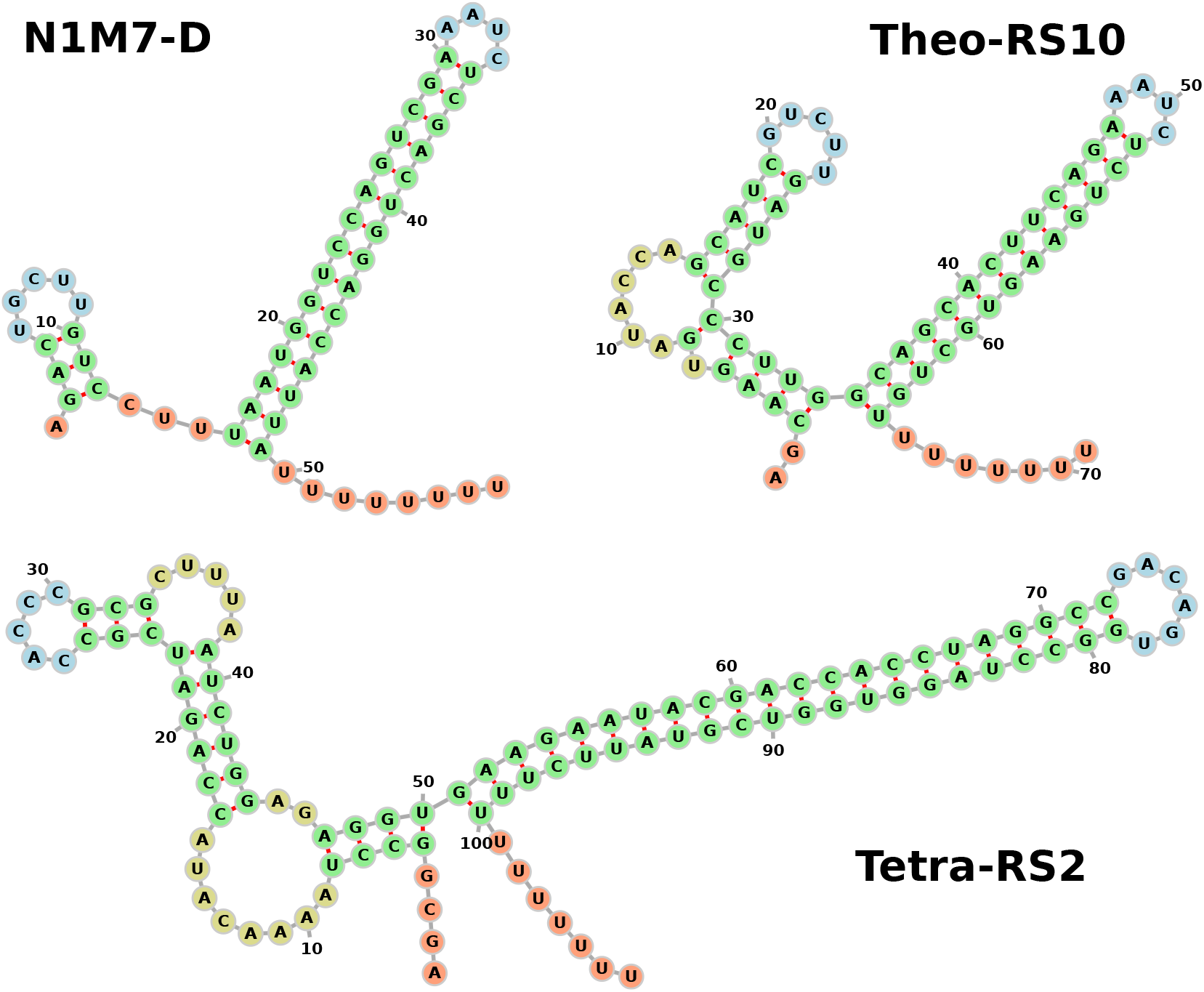
Minimum free energy secondary structures of the neomycin riboswitch N1M7-D, the theophylline riboswitch RS10 [12], and the tetracycline riboswitch RS2 [13] drawn with *forna* [51]. Colors mark different structural features (hairpin loops in blue, interior loops and bulges in light brown, stems in green, and exterior loops in red). All riboswitches display a long terminator hairpin followed by an 8 nt poly-U stretch at their 3’ end. In contrast to the other constructs shown, N1M7-D exhibits a remarkably small structure at its 5’ end.

## B. Supplementary Data

**Table B.1:**
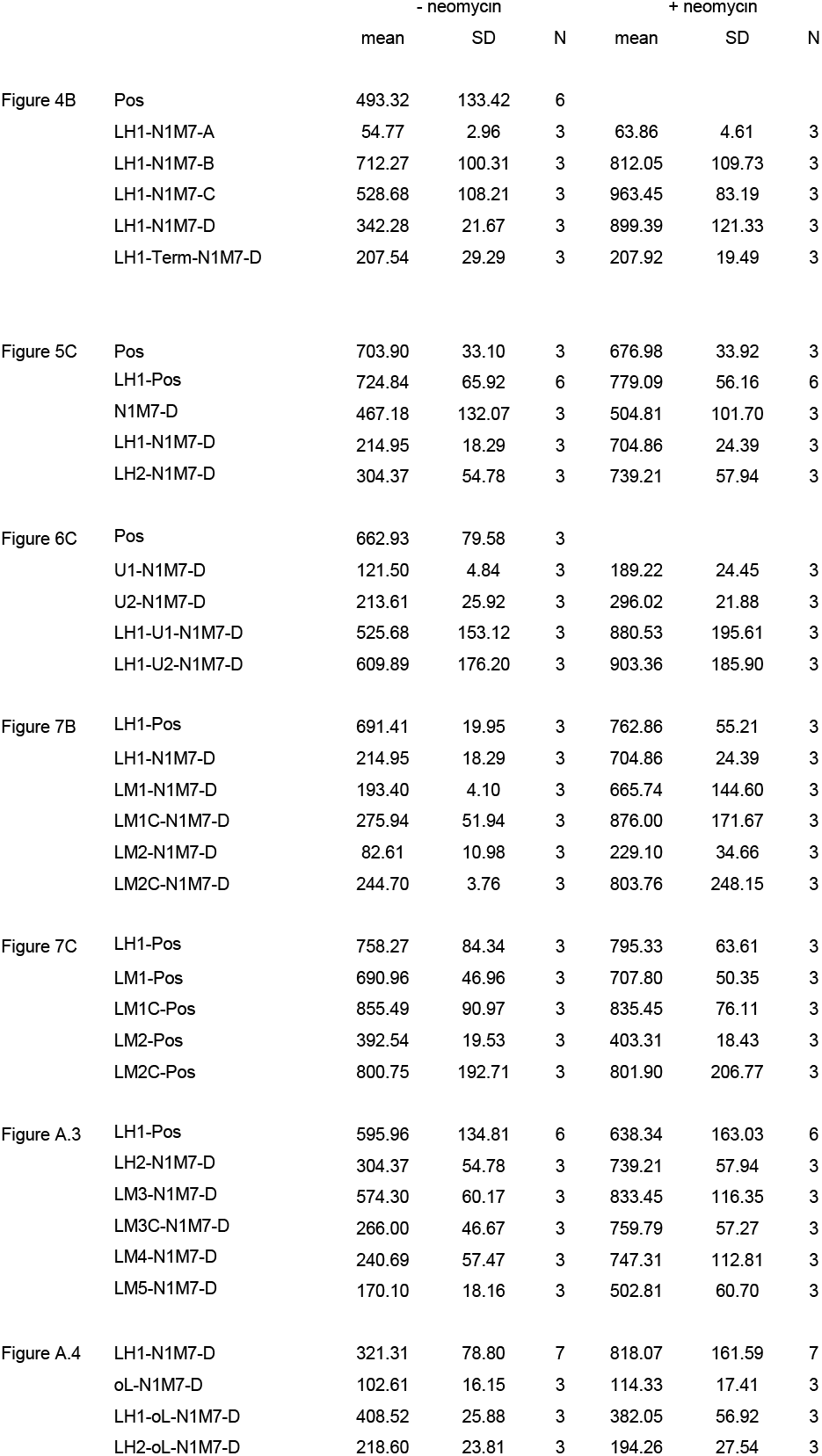
Overview of all constructs analysed in this work and which figures they appear in, along with their corresponding fluorescence intensity data (mean, standard deviation (SD), and the number of repetitions (N)). Sequence data can be found in a supplementary spreadsheet file.

**Table B.2:** Effects of mutations on the stability of leader hairpins LH1 and LH2. The stability is given as minimum free energy (MFE) of the entire (mutated) hairpin. Activities of the resulting constructs are shown in Fig. 7 and Fig. A.3.

| Hairpin | Mutations | MFE (kcal mol<sup>-1</sup>) |
| --- | --- | --- |
| LH1 | none | -15.9 |
| LM1 | G28C, A29U | -10.9 |
| LM1C | G28C, A29U, U6A, U7G | -18.5 |
| LM2 | U21C, C22G, | -9.7 |
| LM2C | U21G, C22G, G13C, G14C | -18.3 |
| LH2 | none | -13.1 |
| LM3 | C17G, G18C | -5.5 |
| LM3C | C17G, G18C, C3G, G4C | -13.9 |
| LM4 | G13C, A14U | -10.1 |
| LM5 | G13C, A14U, G15C | -5.3 |

## Footnotes

1 https://github.com/xileF1337/riboswitch_design

2 5’-GGAGCAATAACTACTTTGTATTTTG-3’

3 5’-GCCATGTGTAATCCCAGCAGCTGTTACAAACTCAAGAAGG-3’

4 5’-TTCTGAATTCGGCATGGGGTCAGGTGG-3’

5 https://www.bioinf.uni-leipzig.de/Software/BarMap_QA/

